# Kinetics of enzymatic mercury methylation at nanomolar concentrations catalyzed by HgcAB

**DOI:** 10.1101/510180

**Authors:** Swapneeta S. Date, Jerry M. Parks, Katherine W. Rush, Judy D. Wall, Stephen W. Ragsdale, Alexander Johs

## Abstract

Methylmercury (MeHg) is a potent neurotoxin that bioaccumulates in fish. MeHg is generated by anaerobic bacteria and archaea possessing the gene pair *hgcAB*. Although bacterial mercury (Hg) methylation has been characterized *in vivo*, the specific role of HgcAB in catalyzing Hg methylation is not well understood. Here we report the kinetics of HgcAB-mediated Hg methylation in cell lysates of *Desulfovibrio desulfuricans* ND132 at nanomolar Hg concentrations. The enzymatic Hg methylation mediated by HgcAB is highly oxygen-sensitive, irreversible, and follows Michaelis-Menten kinetics with an apparent *K_M_* of 3.2 nM and *V*_max_ of 19.7 fmol·min^-1^·mg^-1^ total protein for the substrate Hg(II). Although the abundance of HgcAB in the cell lysates is extremely low, Hg(II) was quantitatively converted to MeHg at subnanomolar substrate concentrations. Supplementation with ATP, methyltetrahydrofolate, or pyruvate did not enhance MeHg production under the experimental conditions. Insight into the kinetics of Hg methylation catalyzed by HgcAB advances our understanding of the complex global Hg cycle.

## Introduction

Mercury (Hg) occurs naturally in the environment and is released in part from natural sources such as geothermal activity and volcanism. However, large quantities of Hg are released as a result of anthropogenic activities, such as mining operations, coal combustion, and other industrial processes (1). In a complex global cycle, Hg is transformed biotically and abiotically among several major forms including elemental Hg^0^, mercuric Hg(II) and methylmercury (MeHg). MeHg is the most prevalent organomercurial and is a potent neurotoxin (2); thus, human exposure to MeHg is a public health concern. MeHg bioaccumulates in the food web and humans are exposed to this neurotoxin through their diet, particularly by consuming Hg-contaminated fish. Certain anaerobic bacteria and archaea are capable of converting Hg to MeHg. We refer to these microorganisms as Hg methylators hereafter. The physiological role of microbial MeHg production is unclear as Hg methylation apparently does not impart resistance to Hg toxicity (3). Interestingly, several Hg methylators have been shown to methylate Hg and demethylate MeHg simultaneously (4). Hg methylation has been studied in environmental samples and in pure cultures of Hg methylators (5–8). However, Hg methylation rates vary significantly with different strains, as well as in the presence of organic matter and/or thiolates (9,10).

Early investigations of Hg methylation in cell extracts of a methanogen suggested that a methyl group could be transferred from methylcobalamin to Hg by both enzymatic and non-enzymatic processes (11–13). Non-enzymatic Hg methylation by methylcobalamin (Eq. 1) is most favorable at pH 4.5 (12,13).

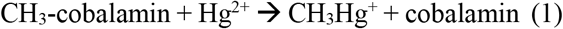

Following the identification and isolation of the sulfate-reducing bacterium *Desulfovibrio desulfuricans* LS as an environmentally relevant Hg methylator (14), a series of studies investigated associations between metabolic pathways, corrinoids, and Hg methylation (15–18). Based on incorporation of ^14^C from radiolabeled precursors into MeHg, the reductive acetyl-coenzyme A (CoA) pathway was implicated with Hg methylation (15,18). High levels of ^14^C incorporation into MeHg were observed in cultures supplemented with L-[3-^14^C]serine and [^14^C]formate (18). In addition, studies with 5-^14^CH_3_-tetrahydrofolate indicated that methyltetrahydrofolate (CH_3_-H_4_folate) may serve as the methyl donor to form MeHg (17). However, enzyme activities associated with the reductive acetyl-CoA pathway were several orders of magnitude lower in *D. desulfuricans* LS compared to acetogens (18). ^57^Co labeling experiments, corrinoid extractions, and mass spectrometry identified cobalamin as the major corrinoid in *D. desulfuricans* LS (16). Additional studies implicated a 40 kDa corrinoid protein in enzymatic Hg methylation, but the specific protein involved was not identified or further characterized (17). As a result of these findings, it was proposed that Hg methylation is a two-step process involving (i) transfer of a methyl group from CH_3_-H_4_folate to a corrinoid protein followed by (ii) transfer of the methyl group from the methylcorrinoid to Hg(II) to form MeHg (17). Later studies suggested that Hg methylation may be independent of the CoA pathway as incomplete-oxidizing sulfate reducing bacteria are able to methylate Hg, but do not use the CoA pathway in major carbon metabolism (19).

The genetic basis of Hg methylation was not well understood until recently (20). The *hgcAB* gene pair was shown to be required for Hg methylation in the sulfate-reducing bacterium *Desulfovibrio desulfuricans* ND132 and the iron-reducing bacterium *Geobacter sulfurreducens* PCA (20). Deletion of either gene resulted in a complete loss of Hg methylation activity. To date, all strains of bacteria and archaea with *hgcAB* genes that have been assayed for Hg methylation are capable of producing MeHg (3,6).

Based on sequence analysis and homology modeling it was predicted that *hgcA* encodes a protein consisting of a corrinoid binding domain (CBD) facing the cytosol and a transmembrane domain (TMD) anchored in the cytoplasmic membrane (Fig. 1A) (20). The CBD of HgcA is homologous to the CBD of the corrinoid iron-sulfur protein (CFeSP) from the reductive acetyl-CoA (Wood-Ljungdahl) pathway of carbon fixation (20), but the TMD has no detectable sequence similarity to any known protein (20). The CBD of HgcA contains a strictly conserved Cys (C93) that is critical for Hg methylation activity *in vivo* (21). The gene *hgcB* almost always appears immediately downstream of *hgcA* and encodes a ferredoxin with two [4Fe-4S] binding motifs (20). HgcB has a unique architecture among ferredoxins, consisting of two [4Fe-4S] cluster-binding motifs (CX_2_CX_2_CX_3_CP), an additional strictly conserved Cys residue (C73), and a pair of conserved cysteines at the C-terminus (C94/95) (Fig. 1A) (20,21). It is not known whether HgcA and HgcB form a multiprotein complex. However, we refer to these proteins collectively as HgcAB hereafter to emphasize that both are required for Hg methylation.

**Figure 1.**
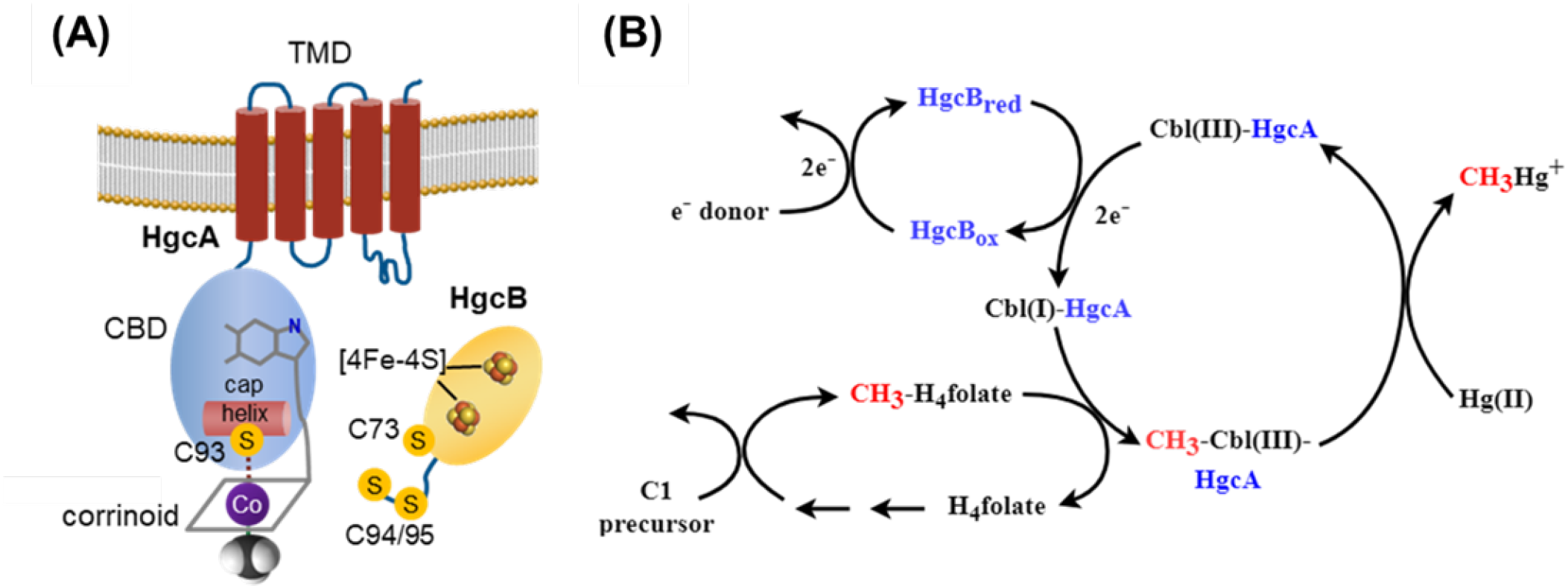
Structural features and proposed mechanism of HgcAB-mediated Hg methylation. (A) Cartoon representation of sequence-based structural models of HgcA and HgcB. Key components and features are labeled. (B) Proposed roles of HgcA and HgcB in Hg methylation.

A general mechanism for Hg methylation by HgcAB is shown (Fig. 1B). The reaction cycle begins with HgcB providing low-potential electrons to the oxidized corrinoid cofactor of HgcA to generate the supernucleophilic Co(I) state. The Co(I)-corrinoid then accepts methyl group from a methyl donor such as CH_3_-H_4_folate to form a CH_3_-Co(III)-corrinoid. The resulting methylcorrinoid then transfers its methyl group to a Hg(II) substrate, either directly or through a ligand exchange reaction (22,23), to form CH_3_Hg^+^. After methyl transfer, the corrinoid is in an oxidized state. Thus, every turnover cycle requires low-potential electrons, donated by HgcB, to reduce the corrinoid to the Co(I) state. How the Hg(II) substrate enters cells and interacts with HgcAB is currently unknown, but uptake and transport of Hg(II) into the cytoplasm are essential and may limit methylation rates observed *in vivo* (24,25).

To evaluate the kinetics of Hg methylation mediated by HgcA and HgcB independent of transport processes, we performed Hg methylation assays in cell lysates of *D. desulfuricans* ND132 (ND132). ND132 is a sulfate-reducing bacterium that is often used as a model organism for studying Hg methylation due to robust Hg methylation by ND132 and similarity to *D. desulfuricans* LS (3,26). The present study using ND132 builds on the foundation of earlier studies conducted with the strain *D. desulfuricans* LS, which was not sequenced and is considered lost (15–18). Owing to advances in MeHg analysis, we perform experiments at much lower and more environmentally relevant levels of Hg (0.5 to 60 nM compared to 0.5 to 8 mM previously used for studies with *D. desulfuricans* LS (17)). We follow Hg methylation rates in ND132 cell lysates over time and study the effects of pH, temperature and total protein concentration. We determine initial rates of Hg methylation in cell lysates as a function of Hg(II) concentration and calculate apparent kinetic parameters (*K_M_* and *V*_max_) for ND132 wild-type (WT) and compare results to an *hgcAB* deletion mutant *(ΔhgcAB)*. Furthermore, we evaluate the contributions of enzymatic Hg methylation, non-enzymatic Hg methylation and whether the Hg methylation reaction is reversible or a separate MeHg demethylation pathway exists in ND132. We also conduct a set of targeted experiments to gain insights into aspects of the proposed mechanism of Hg methylation by HgcA and HgcB. Specifically, we investigate the sensitivity of Hg methylation to oxygen exposure, evaluate a potential role of ATP on Hg methylation and determine the effect of supplementation of the cellular metabolites CH_3_-H_4_folate and pyruvate, which are proposed methyl and electron donors, respectively.

## Results

### Time-dependence of Hg Methylation and MeHg Demethylation

The time course for the conversion of Hg(II) to MeHg in cell lysates of ND132 WT and *ΔhgcAB* was determined over a period of up to 92 h. The levels of MeHg produced in WT cell lysates increased at a rate of 1.31 ± 0.14 nM/h within the first 2 h, decreasing over time and plateauing between 24 and 92 h (**Fig. 2A**). At the 92-h time point, 56.4% of the added Hg(II) was converted to MeHg. No significant MeHg production was observed in *ΔhgcAB* cell lysates over the course of the experiment. MeHg levels in *ΔhgcAB* cell lysates were < 0.5% of MeHg in WT cell lysates at 92 h, confirming that Hg methylation in ND132 cell lysates is strictly an HgcAB-dependent process. In addition, the lack of significant MeHg production in *ΔhgcAB* cell lysates indicates that non-enzymatic methylation mediated by methylcorrinoids, either free (Eq. 1) or associated with other proteins, did not contribute significantly to MeHg formation in the cell lysates. Initial rates of Hg methylation measured over the first 2 h after Hg(II) addition were used for subsequent experiments.

**Figure 2.**
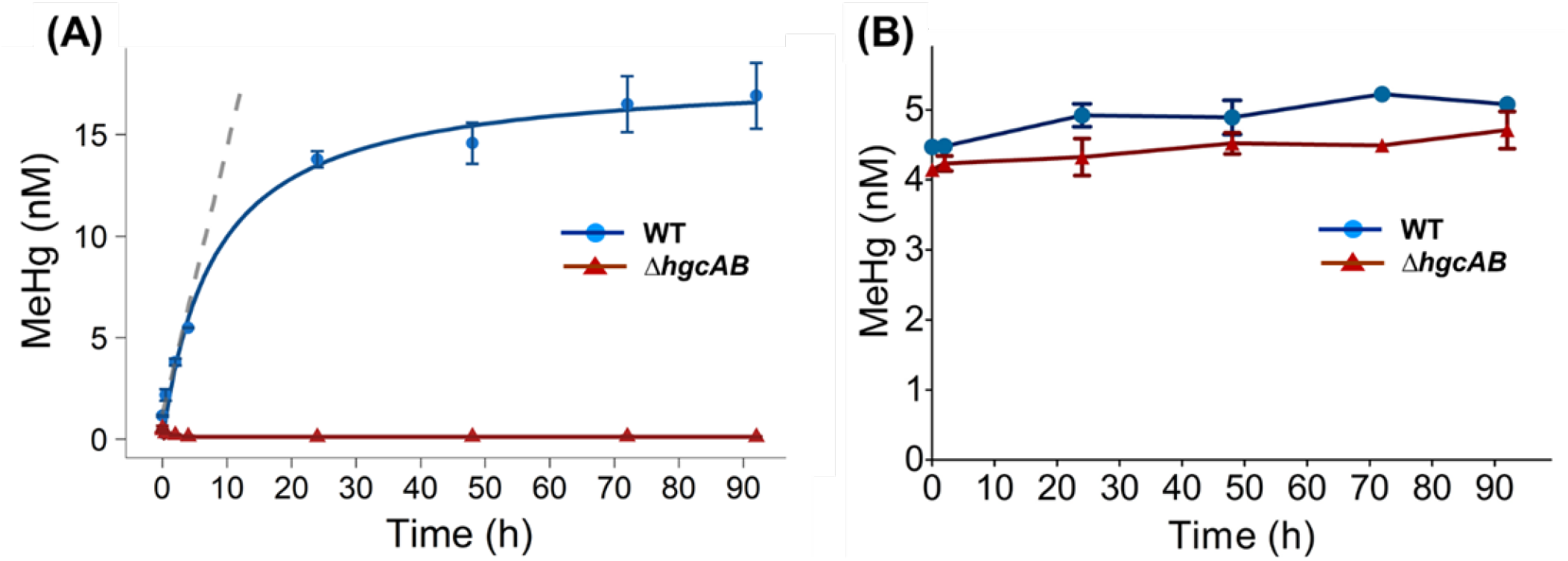
Time dependence of Hg methylation and MeHg demethylation in ND132. (A) Hg methylation in cell lysates of ND132 WT (blue) and *ΔhgcAB* (red) (1.5 mg/mL total protein concentration) under strictly anaerobic conditions at 32 °C in the presence of 30 nM Hg(II). The blue line shows a nonlinear fit of the concentration data and the gray dashed line shows a linear fit (R^2^ = 0.98) for time points between 0 and 2 h to determine initial rates. (B) MeHg concentrations in ND132 cell lysates of WT (blue) and *ΔhgcAB* (red) as a function of time under a similar experimental setup as (A) but with 5 nM MeHg as the substrate. Error bars represent standard deviation between duplicate set of samples (N=2).

To determine whether a fraction of Hg was reduced to Hg^0^ over the course of the experiment and to close the Hg mass balance, we also measured the total Hg (THg) for every time-point. No significant changes in THg levels were observed, eliminating the possibility that Hg was lost through reduction to Hg^0^ over the course of the experiment (**Fig. S1**).

To determine whether MeHg is demethylated in ND132 cell lysates, we incubated WT cell lysates with 5 nM MeHg and measured MeHg levels over time up to 92 hours. Considering that demethylation may be independent of HgcAB, we compared the demethylation of MeHg in cell lysates of both ND132 WT and the *ΔhgcAB* mutant. No demethylation was observed for either strain (**Fig. 2B**). Therefore, we conclude that Hg methylation by HgcAB is irreversible and that there is no other mechanism independent of HgcAB for MeHg demethylation in ND132 cell lysates.

### pH, temperature, and total protein concentration dependence of Hg Methylation

To provide further evidence that Hg methylation in ND132 cell lysates is an enzymatic process, we determined Hg methylation rates as a function of pH, temperature, and total protein concentration. The rate of MeHg production in WT cell lysates increased by a factor of 6.5 between pH 4.0 (0.32 ± 0.06 nM/h) and pH 8.0 (2.12 ± 0.03 nM/h) (**Fig. 3A**). This result is in contrast to the pH-rate profile for the non-enzymatic methylation of Hg(II) by methylcobalamin (Eq. 1) in which the highest Hg methylation rates were observed at pH 4.5, and decreased significantly at pH 6.0 and higher (13). Hg methylation rates in WT cell lysates varied by an order of magnitude between 4 °C (0.22 ± 0.00 nM/h) and 32 °C (1.91 ± 0.12 nM/h), and between 32 °C and 50 °C (0.14 ± 0.01 nM/h), with a fairly narrow optimum centered around 32 °C (**Fig. 3B**). The observed temperature dependence is consistent with the optimal growth temperature of ~32 °C for ND132 cultures and a lack of any substantial growth at 45 °C and above (3). The observed pH and temperature dependencies suggest that MeHg is predominantly formed by an enzymatic process dependent on HgcAB under the experimental conditions.

**Fig. 3.**
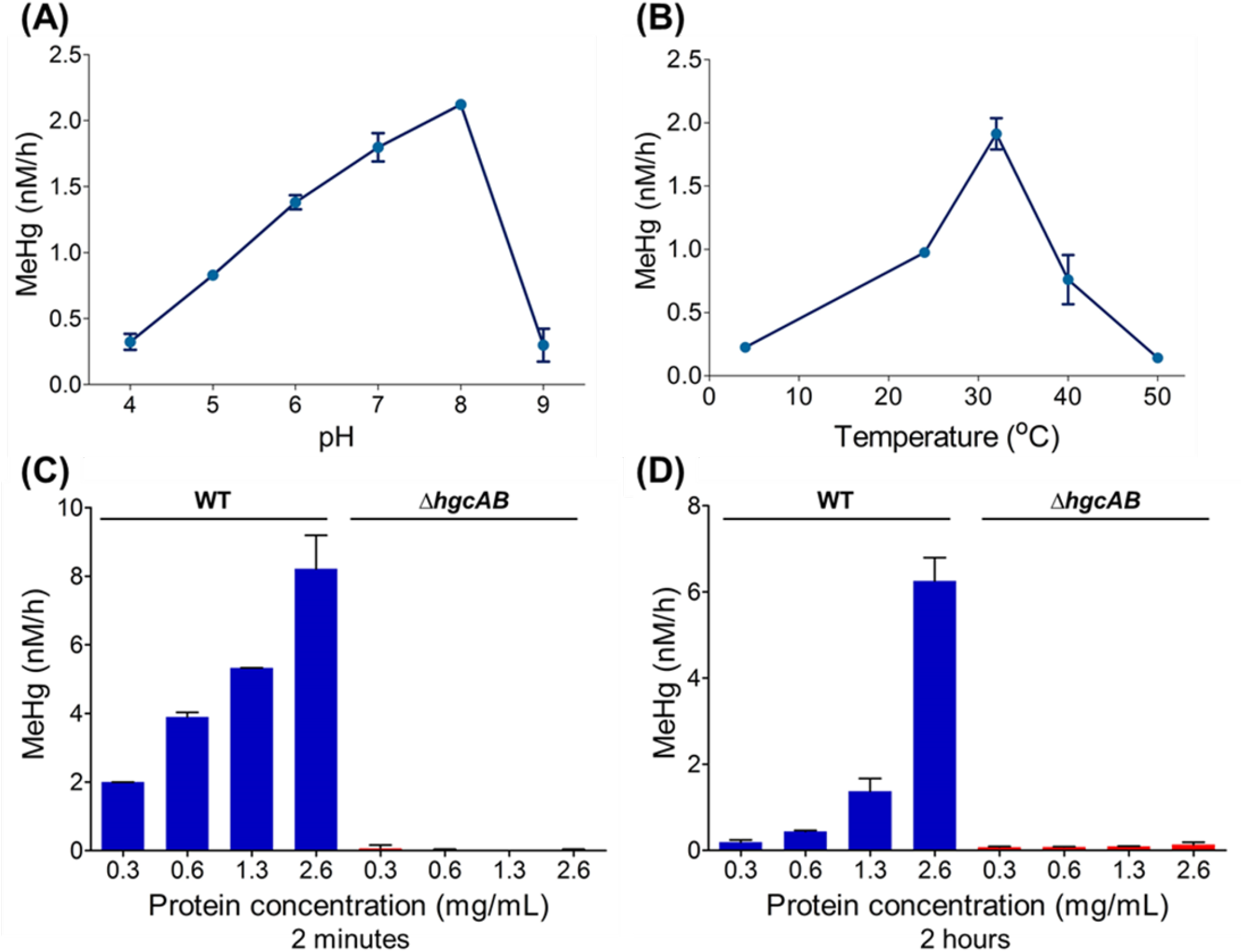
Dependence of Hg methylation rates on pH, temperature and total protein concentration. Hg methylation rates in cell lysates of ND132 WT measured as a function of (A) pH (4, 5, 6, 7, 8, and 9), (B) temperature (4, 24 (RT), 32, 40, and 50 °C), and (C and D) total protein concentration. Samples were incubated with 30 nM Hg(II) and harvested for MeHg analysis at 2 h, except in (C) where samples were harvested at 2 minutes after Hg(II) addition. pH 7, 32 °C, and strictly anaerobic conditions correspond to standard experimental conditions. Error bars represent standard deviation between duplicate set of samples (N=2).

To investigate the effect of total protein concentration on the rate of Hg methylation in cell lysates, we measured MeHg production rates at 2 min (**Fig. 3C**) and 2 h (**Fig. 3D**) over a range up to ~8 times the initial total protein concentration (0.3 mg/mL to 2.6 mg/mL). The highest total protein concentration achievable is limited by the volume of buffer required to resuspend cell pellets prior to cell lysis. As expected, the rate of MeHg formation increased with increasing concentrations of total protein. Although the initial rates determined within 2 min of the addition of Hg(II) were higher and increased linearly in response to total protein concentration, methylation rates determined at 2 h increased exponentially with increasing total protein concentrations.

### Enzyme kinetics of HgcAB-mediated Hg methylation

To examine the effects of increasing substrate concentrations on the rate of MeHg production, we incubated ND132 cell lysates (1.5 mg/mL total protein concentration) with HgCl_2_ as a substrate at concentrations ranging from 0.5 nM to 60 nM. *ΔhgcAB* cell lysates were used as a negative control. Initial MeHg formation rates followed Michaelis-Menten kinetics with a maximum rate *V*_max_ = 19.7 (± 0.35) fmol·mg total protein^-1^ min^-1^ and *K_M_* = 3.2 nM (± 0.26) for Hg(II) (**Fig. 4A**). CH_3_-H_4_folate is the presumed methyl donor for MeHg. As described later, CH_3_-H_4_folate levels were not rate-limiting under the current experimental conditions. At low concentrations of Hg(II), 0.5 nM and 1 nM, nearly 100% added Hg(II) was converted to MeHg (**Fig. 4B**).

**Fig. 4.**
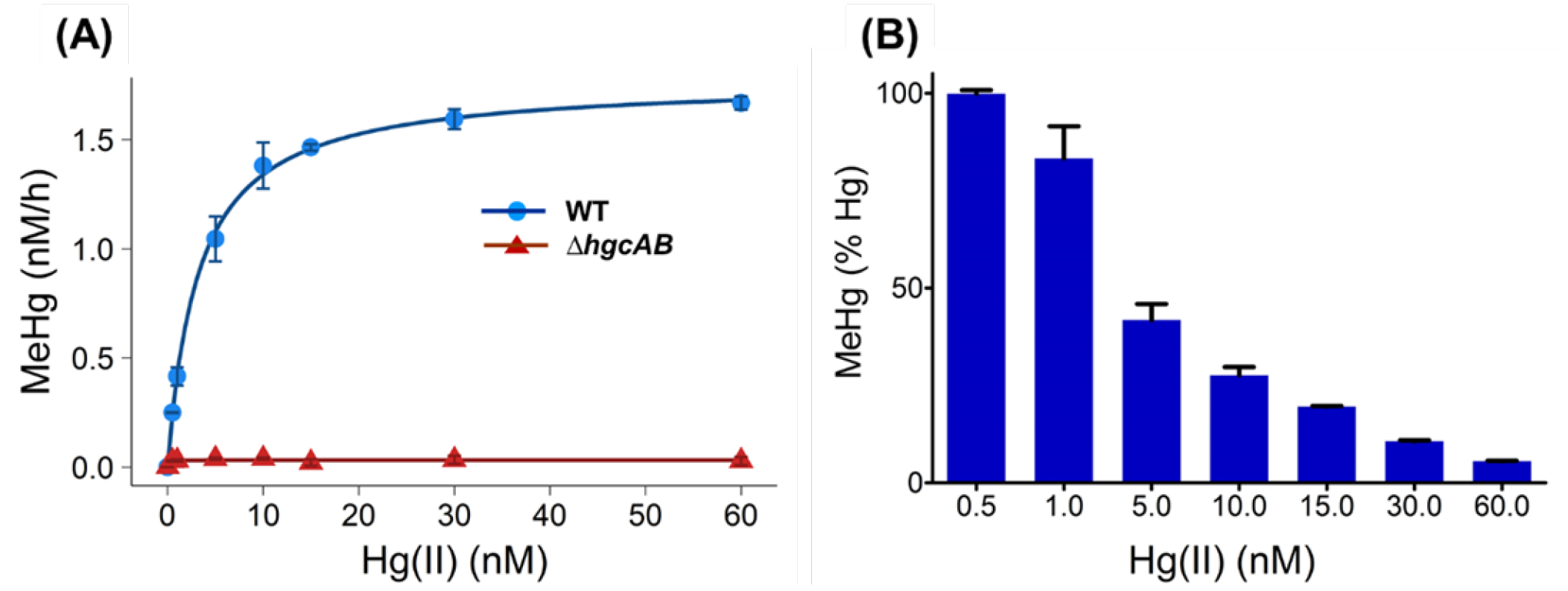
Dependence of Hg methylation rates on Hg substrate concentration. (A) Initial rates of Hg methylation in ND132 cell lysates (1.5 mg/mL total protein) in response to increasing concentrations of added Hg(II) from 0.5 nM to 60 nM at 32 °C for 2 h under strictly anaerobic conditions for WT (blue circles) and *ΔhgcAB* strains (red triangles). Hg methylation rates in WT cell lysates were fitted to the Michaelis-Menten equation with *V*_max_ = 1.77 ± 0.03 nM/h and *K_M_* = 3.2 ± 0.26 nM (blue line). (B) Percentage of Hg(II) converted to MeHg within 2 h as a function of initial Hg(II) concentration. Error bars represent standard deviation between duplicate set of samples (N=2).

RT-PCR data from ND132 (20) and proteomics studies in ND132 (27) and *G. sulfurreducens* PCA (28) indicated that the abundance of HgcA and HgcB in Hg methylators is extremely low. Owing to the low abundance and resultant difficulties in determining accurate concentrations of HgcA and HgcB in the cell lysates, we used the obtained kinetic parameters to estimate turnover numbers (*k*_cat_), catalytic efficiencies *(k_cat_/K_M_)*, and free energies of activation based on transition state theory (29,30) for a range of enzyme concentrations expressed as a fraction of total cellular protein concentration (**Table 1**, **Fig. S2**). It should be noted that these values are calculated solely for the purpose of gaining further insights into the functioning of HgcAB based on estimated kinetic parameters. Additionally, since the value for *V*_max_ reported here is normalized to the total protein in the lysates, the *V*_max_ of the pure enzyme is expected to be higher. Nevertheless, the value of *K_M_*, which is independent of the enzyme concentration, suggests that HgcA and HgcB readily bind Hg(II) at very low concentrations.

**Table 1.** Turnover numbers (k_cat_), catalytic efficiencies *(k_cat_/K_M_)* and activation free energies (AG^ŧ^) of enzymatic Hg methylation for a range of postulated enzyme (HgcA) concentrations expressed as a fraction of total cell protein.

| **[E] (% of total protein)** | **$k_{\text{cat}}$ (s<sup>-1</sup>)** | **$k_{\text{cat}}/K_M$ (M<sup>-1</sup>·s<sup>-1</sup>)** | **$\Delta G^\ddagger$ (kcal·mol<sup>-1</sup>)** |
| --- | --- | --- | --- |
| 0.1 | $1.2 \cdot 10^{-5}$ | $4 \cdot 10^3$ | 24.1 |
| 0.01 | $1.2 \cdot 10^{-4}$ | $4 \cdot 10^4$ | 22.8 |
| 0.001 | $1.2 \cdot 10^{-3}$ | $4 \cdot 10^5$ | 21.4 |
| 0.0001 | $1.2 \cdot 10^{-2}$ | $4 \cdot 10^6$ | 20.1 |

### Effect of oxygen and cellular metabolites of Hg methylation

Based on the current mechanistic hypothesis, each MeHg formation turnover cycle requires the transfer of low-potential electrons to reduce the corrinoid of HgcA to the Co(I) state to accept a methyl group from a methyl donor (**Fig. 1B**). The presence of oxygen would raise the redox potential in solution and rapidly oxidize any Co(I) due to the low midpoint potential of the Co(I)/Co(II) couple (31,32). Therefore, corrinoid-dependent methyl transfer reactions are highly redox-sensitive. Exposure to oxygen would interrupt the transfer of methyl groups to the corrinoid and inhibit MeHg formation. To test the effect of oxygen on Hg methylation, MeHg production in ND132 cell lysates was measured under both aerobic (exposed to ambient oxygen levels) and anaerobic conditions (standard experimental conditions in the glove box under N_2_ and < 0.6 ppm O_2_). MeHg formation was inhibited by 95% (P < 0.001) in the presence of ambient oxygen (**Fig. 5A**). As expected, no effect of ambient oxygen on MeHg formation was observed in the *ΔhgcAB* cell lysates. Thus, the observed oxygen sensitivity of HgcAB-mediated Hg methylation in cell lysates of WT ND132 is consistent with the proposed requirement for low-potential electrons (i.e., < −450 mV) for the corrinoid in HgcA to achieve the Co(I) state (31,32). Additionally, the presence of oxygen may also affect the oxidation state of the two [4Fe-4S] clusters in HgcB.

**Fig. 5.**
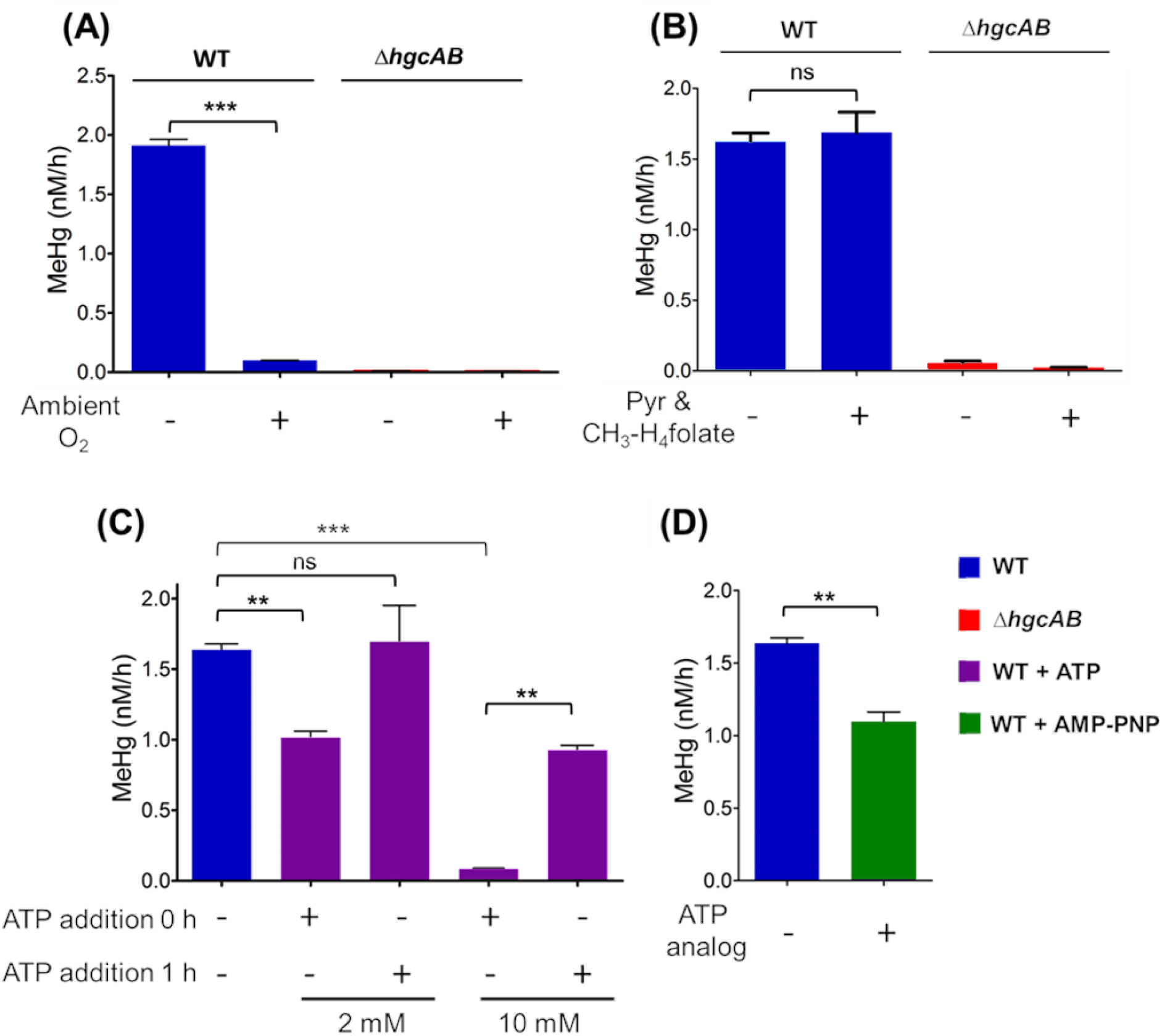
Effect of ambient oxygen and key cellular metabolites on Hg methylation. (A) Hg methylation rates in WT ND132 cell lysates and *ΔhgcAB* under aerobic (exposed to ambient oxygen) and anaerobic conditions (N_2_ environment, < 0.6 ppm O_2_). (B) Hg methylation rates in WT ND132 cell lysates and *ΔhgcAB* under unamended conditions compared to the Hg methylation rates for samples amended with 16.8 μM CH_3_-H_4_folate and 10 mM pyruvate (Pyr). (C) Effect of 2 mM ATP and 10 mM ATP on the concentration of MeHg produced at 2 h in the presence of 30 nM Hg(II) compared to the unamended sample. ATP was supplemented before the addition of Hg(II) (ATP addition 0 h) or 1 h after the addition of Hg(II) (ATP addition 1 h). (D) Effect of 5 mM non-hydrolyzable ATP analog AMP-PNP on MeHg production compared to the unamended sample. Data were analyzed with a two-sample t-test (ns = P > 0.05, ** = P ≤ 0.01, *** = P ≤ 0.001). Error bars represent the standard deviation between duplicates (N=2).

CH_3_-H_4_folate was previously proposed as a potential methyl donor required for the formation of MeHg. Furthermore, no Hg methylation was detected in cell lysates of *D. desulfuricans* LS without the addition of 10 mM pyruvate, wherein pyruvate was proposed to play a role in the generation of reductant such as reduced ferredoxin (15,17,18). To determine whether the exogenous addition of 16.8 μM CH_3_-H_4_folate and 10 mM pyruvate, as used in previous studies, leads to increased Hg methylation, we compared the Hg methylation rates under our standard experimental conditions (i.e., unamended) to that of samples amended with pyruvate and CH_3_-H_4_folate. No significant changes in the rate of MeHg formation were observed in response to the combined addition of 16.8 μM CH_3_-H_4_folate and 10 mM pyruvate (P > 0. 05) (**Fig. 5B**). Because the Hg(II) concentration used for the current experiments (30 nM) is much lower than the levels used for assays with *D. desulfuricans* LS (0.7 mM), we propose that the availability of pyruvate and CH_3_-H_4_folate were not limiting factors during MeHg formation under our experimental conditions. In addition, involvement of a physiological methyl donor other than CH_3_-H_4_folate cannot be ruled out.

Previous studies have proposed a role for ATP driving the uptake or methylation of Hg (24). To determine whether Hg methylation in cell lysates is ATP-dependent, we measured the effect of exogenously added ATP on MeHg formation independent of uptake processes. ATP was added to cell lysates at a concentration of 2 mM, which is close to the average physiological ATP concentration determined for *Escherichia coli* cells (33), and in excess, 10 mM. ATP was added at 0 h, i. e., before adding Hg. In a separate set of samples, ATP was supplemented 1 h after Hg was added (ATP at 1 h). Samples were harvested 2 h after the addition of Hg(II) in both cases. In selecting the timing of ATP supplementation, we assumed that a substantial amount of endogenous ATP present in the lysate would decrease after 1 h, and that the addition of exogenous ATP at that time point would increase MeHg production if Hg methylation is ATP-dependent. The 1 h time point is within the initial, linear rate period of the Hg methylation progress curve (**Fig. 2**). Interestingly, MeHg production decreased significantly (P < 0.05) by 38% and 95% respectively, upon addition of exogenous ATP at concentrations of 2 mM and 10 mM compared to unamended samples (**Fig. 5C**). Addition of 2 mM ATP at 1 h did not significantly increase MeHg production. However, addition of 10 mM ATP after 1 h significantly decreased the total amount of MeHg formed after 2 h. The amount of MeHg produced after supplementation of 10 mM ATP was about 56% of that formed under unamended conditions.

To rule out the possibility that the decrease in MeHg production in the presence of ATP may have arisen from the loss of Hg(II) through ATP-dependent reduction to Hg(0), we determined the concentration of THg in ND132 cell lysates after 2 h. No significant loss in THg was observed in samples with or without added ATP (**Fig. S3**), indicating that the addition of ATP did not result in a loss of Hg(II) through reduction to Hg(0).

Genomic analysis of Hg methylators indicated the presence of a gene encoding a reductive activator of corrinoid proteins (RACo) in several Hg methylators (**Table S1, Fig. S3**). RACo harbors an ATP-binding site and has been shown to form a complex with CFeSP (34). Methylation of CFeSP occurs only in the Co(I) state, which can be oxidized sporadically to the inactive Co(II) state every 100-2,000 turnovers (35). RACo serves as the activator of inactive Co(II)-CFeSP to active Co(I)-CFeSP by coupling the hydrolysis of ATP to the reduction of Co(II) to Co(I) (36). In *Moorella thermoacetica* and *Carboxydothermus hydrogenoformans*, the gene encoding RACo is located in close proximity to the genes encoding enzymes of the reductive acetyl-CoA (Wood-Ljungdahl) pathway (35). Although *D. desulfuricans* ND132 lacks CFeSP and does not have a complete Wood-Ljungdahl pathway (20), we hypothesize that a RACo homolog, commonly present among many methylators, may serve a similar ATP-dependent corrinoid activation role for HgcA.

If MeHg formation is dependent on energy derived from ATP hydrolysis, such as through ATP-dependent corrinoid activation by RACo, competitive inhibition of ATP-hydrolysis sites by a non-hydrolyzable ATP analog would be expected to decrease MeHg production. Therefore, we evaluated the effect of the addition of the non-hydrolyzable ATP analog adenylyl-imidodiphosphate (AMP-PNP) on MeHg formation in ND132 cell lysates. AMP-PNP was added at a concentration of 5 mM, an amount that is expected to compete with endogenous ATP for ATP hydrolysis sites of most ATPases (37). The addition of AMP-PNP resulted in a decrease of MeHg production by 33% (P < 0.05) compared to the unamended control (**Fig. 5D**). This effect on MeHg production is similar to the decrease observed after addition of ATP. However, the mechanism by which both ATP and AMP-PNP decrease MeHg production is unclear.

## Discussion

Why bacteria and archaea methylate mercury remains a mystery, as methylation apparently does not impart these organisms with mercury resistance (3). Microbial Hg methylation has been the subject of investigation since the late 1960s (11,38). The discovery of *hgcAB* offered a first glimpse at the unique bioinorganic chemistry involved in the methylation of Hg. In this study, we investigated the cellular Hg methylation mediated by HgcAB. We measured the contributions of enzymatic and non-enzymatic processes to Hg methylation in cell lysates of anaerobic model Hg methylator *D. desulfuricans* ND132. With the aim of deciphering HgcAB mediated molecular mechanism of Hg methylation, we tested the effect of various biochemical parameters on Hg methylation. The mechanistic hypothesis of Hg methylation (**Fig. 1B**) was developed based on the sequences of HgcA and HgcB, as well as early investigations of Hg methylation in *D. desulfuricans* LS (15–18).

As expected, methylation activity was detected in lysates of the WT, but not in lysates of the *ΔhgcAB* strain confirming that HgcA and HgcB are essential for converting inorganic Hg(II) to MeHg (**Fig. 2A**). It should be noted that ND132 and other strains of sulfate-reducing bacteria have been reported to simultaneously methylate Hg(II) and demethylate MeHg (3,4). In our experiments with ND132 cell lysates, however, we did not observe demethylation of added MeHg over a period of several days (**Fig. 2B**). Hg methylation in ND132 under the tested conditions is, therefore, irreversible and does not affect the Hg methylation rate parameters.

At the lowest lysate concentration (0.3 mg/mL protein), the initial rate of MeHg formation determined over a period of two minutes after Hg(II) addition was higher than the rate determined over a two-hour period. However, the rate increased linearly with the concentration of the lysate in the two-minute experiments, while it increased exponentially with lysate concentration in the two-hour experiments (**Fig. 3C and D**). The linear increase in the two-minute experiments indicates that the rates of MeHg formation were limited only by the overall enzyme concentration. However, the exponential increase of Hg methylation rate at higher lysate concentrations in the two-hour experiments suggests that the rate is also dependent on the concentration of a secondary substrate present in the lysates, which may have been depleted over time. While this secondary substrate is currently unknown, our findings are consistent with previous studies with *D. desulfuricans* LS in which the rate of MeHg formation also increased exponentially with increasing cell lysate concentrations, suggesting that Hg methylation involves two or more components, such as a methyl donor, electron donor, and Hg(II) substrate (17). Nevertheless, Hg(II) substrate concentration dependence under the conditions tested in our experiments followed Michaelis-Menten kinetics (**Fig. 4A**). Although the estimated turnover numbers (k_cat_ ~ 1 · 10^-5^ to 1·10^-2^ s^-1^) are very low compared to an average enzyme (39) (**Table 1**), the low *K_M_* of 3.2 nM yields catalytic efficiencies (*k*_cat_/*K_M_* ~ 4·10^3^ – 4·10^6^ M^-1^ s^-1^) that fall within reasonable ranges.

Hg(II) is toxic to many enzymes as a result of its high affinity for biological thiolates (SR^-^) with stability constants (log β) for Hg(II)-bis-thiolate [Hg(SR)_2_] complexes of up to 45 (40). Hg(II) added to cell lysates as HgCl_2_ at nanomolar concentrations is therefore expected to form highly stable complexes with intracellular thiolates. Although the thermodynamic stability of Hg(SR_)2_ species is extremely high, ligand exchange of such complexes with other free thiolates is rapid (41). Therefore, the low *K_M_* and quantitative conversion of Hg(II) to MeHg at low substrate concentrations (**Fig. 4B**) are remarkable and suggest that the active site may use thiolate ligands to bind Hg(II) by a ligand exchange mechanism, similar to mechanisms previously described for MerA and MerB (42,43). A series of conserved cysteines identified in sequences of HgcB (C94, C95 and C73) may facilitate such ligand exchange processes. Site-directed mutagenesis experiments in ND132 showed that a C73A mutation eliminates Hg methylation in vivo, but only one of the two C-terminal cysteines (C94, C95) is essential for activity (21).

Based on the hypothesized roles for HgcA and HgcB (**Fig. 1B**), in addition to a Hg(II) substrate, Hg methylation requires both a source of electrons to regenerate the Co(I) state of the corrinoid cofactor and a methyl donor. Although the source of electrons is unknown, HgcB likely assumes a key role in shuttling electrons from an unidentified electron donor to the corrinoid cofactor of HgcA. If the intracellular molar ratio of HgcA to HgcB remains constant, the generation of electrons may become rate-limiting as it depends on active metabolic processes, which are expected to decrease in the cell lysates over time. Furthermore, the immediate methyl donor to HgcA remains unknown, as supplementation of CH_3_-H_4_folate did not result in an increased rate of Hg methylation over a period of two hours (**Fig. 5B**).

Experiments to evaluate the role of ATP-driven processes showed that supplementation of ATP leads to a substantial decrease in Hg methylation activity in cell lysates (**Fig. 5C**). Therefore, increasing ATP concentrations does not enhance intracellular Hg methylation. A similar decrease in Hg methylation was observed with the competitive nonhydrolyzable analog AMP-PNP (**Fig. 5D**). Thus, we could not confirm the hypothesis that reductive activation by RACo or a similar ATP-dependent activator is essential for Hg methylation. Alternatively, ATP may affect Hg speciation or impact other, yet to be identified, cellular components related to Hg methylation.

Several key observations in this study are consistent with results obtained from studies with *D. desulfuricans* LS. The majority (> 95%) of cellular Co in *D. desulfuricans* LS was reported to be associated with macromolecules (16). In ND132 cell lysates, no non-enzymatic Hg methylation was observed and, in the *ΔhgcAB* mutant, Hg methylation was negligible, indicating that HgcAB is indeed responsible for all Hg methylation. A 40 kDa protein, being the only major Co-containing protein in *D. desulfuricans* LS (17), agrees with the molecular mass of ND132 HgcA (36.7 kDa without its corrinoid cofactor) or with a HgcAB complex (1:1 molar ratio) (~49 kDa), although it is not known whether HgcA and HgcB form a stable complex.

However, there are several differences between the experimental conditions used in the present study and those performed previously with *D. desulfuricans* LS. While the typical concentration of Hg(II) in our experiments was 30 nM, which is similar to levels used for methylation experiments with whole cells (25), experiments with *D. desulfuricans* LS were conducted at Hg(II) levels in the mM range (0.5 to 8 mM) (17). Hg toxicity to ND132 cells was apparent through decreased cell density at Hg concentrations above ~ 5 μM (3). Detection limits for the analysis of MeHg by ICP-MS following the current standard EPA methods are in the low pM range (44). The vastly improved detection limits for MeHg enable our studies at significantly lower Hg levels, which are more physiologically relevant and dramatically reduce potential impacts of Hg toxicity to cellular proteins. These advances also allowed us to evaluate Hg methylation kinetics in lysates without supplementing essential metabolites. For example, no Hg methylation was observed with *D. desulfuricans* LS cell extracts unless 10 mM pyruvate and 16.8 μM CH_3_-H_4_folate were added (18). Under our experimental conditions, we observed no effect on Hg methylation activity in ND132 cell lysates of adding an exogenous methyl donor (CH_3_-H_4_folate) or electron source (pyruvate). These differences in the experimental design are reflected in differences between the *V_max_* and *K_M_* values previously determined for *D. desulfuricans* LS (*V*_max_ = 0.728 nmol min^-1^ · mg protein^-1^ and *K_M_* = 0.872 mM) and those described here (*V*_max_ = 19.7 fmol · min^-1^ · mg total protein^-1^ and *K_M_* = 3.2 nM). Differences in the inhibition of methylation activity by ambient oxygen (45% for *D. desulfuricans* LS (17), 95% in this study) suggest that non-enzymatic methylation may have been more prominent in studies with *D. desulfuricans* LS and may have contributed to overall MeHg production. Here we showed that essentially all of the MeHg formed in ND132 is a product of enzymatic activity mediated by HgcAB.

Inhibition of Hg methylation by oxygen is consistent with the proposed HgcAB-mediated turnover cycle (**Fig. 1B**). Exposure of cell lysates to ambient oxygen could inactivate either FeS-containing HgcB or corrinoid-containing HgcA or oxidize the active Co(I)-corrinoid to Co(II) or Co(III). As a consequence, in the presence of oxygen HgcA would not be able to accept a methyl group, thus interrupting the turnover cycle.

We speculate that the fraction of HgcA is equivalent to approximately 0.0004% of the total protein concentration in cell lysates (see Supplementary Materials), which would correspond to approximately 1 in 220,000 proteins in the cell lysate. Such low abundance would explain difficulties in identifying HgcA protein fragments and low transcript levels in previously published studies (20,27,28). The calculated kinetic parameters resulting from this HgcA abundance estimate are *k_cat_* = 3·10^-3^ s^-1^, and *k_cat_/K_M_* = 9·10^5^ M^-1^ · s^-1^. Consequently, we speculate that the specific activity of HgcA is close to 4 nmol·min^-1^·mg^-1^ HgcA. Comparing the current estimated rate constant for the HgcAB-mediated enzymatic Hg methylation to previously published non-enzymatic rate constants for methylcobalamin, the enzymatic reaction is approximately 2,500 times faster at pH 7.0 than the non-enzymatic reaction, which is fastest at pH 4.5 (12,13). Although the cofactor-replete expression and purification of HgcA and HgcB presents significant challenges, studies with purified HgcA and HgcB will be essential to validate the postulated kinetic parameters.

In conclusion, we demonstrate that cell lysates containing HgcA and HgcB exhibit Hg methylation activity even at sub-nanomolar Hg(II) concentrations, which is remarkable considering Hg(II) is known to form thermodynamically stable complexes with the abundant pool of cellular thiols present in cells. Future studies aimed at characterizing the molecular structures, and functions of HgcA and HgcB are essential to delineate the interplay between methyl donor, electron donor, and the Hg(II) substrate in cellular Hg methylation. Taking into consideration that deletion of *hgcAB* does not significantly alter the growth of Hg methylators, understanding the cellular biochemistry of Hg methylation could help inform strategies to limit MeHg formation in Hg methylating bacteria and archaea, and thus curtail the formation of this potent neurotoxin in the environment.

## Experimental procedures

### Strains and culture conditions

*Desulfovibrio desulfuricans ND132* cells (wild-type (WT) or *ΔhgcAB* mutant) were grown in modified MOY medium containing 40 mM fumarate, 40 mM pyruvate, and 1.2 mM thioglycolate and 1 mM cysteine HCl as reducing agents for 3 days (OD_600_ = 0.3) at 32°C under strictly anaerobic conditions (< 0.6 ppm O_2_) in 2 L media bottles capped with butyl rubber stoppers (21). For the preparation of ND132 cell lysates, all of the following steps were carried out under strictly anaerobic conditions inside a LABmaster Pro glove box (MBraun, Stratham, NH) under an N_2_ atmosphere maintained below 0.6 ppm O_2_. The cultures were harvested at 4 °C by centrifugation for 20 min at 7,500 × *g* in centrifuge bottles with sealing closures. Pellets were washed 2 times with deoxygenated phosphate buffered saline (PBS), pH 7.0. Cell pellets were resuspended in cold resuspension solution (Pierce™ EDTA-free protease inhibitor, Thermo Fisher Scientific, Waltham, MA, Benzonase^®^ Nuclease, MilliporeSigma, St. Louis, MO, and 2 mM dithiothreitol (DTT) in PBS, pH 7.0) and disrupted on ice by four cycles of sonication (50 sec, 70% power) with a Sonic Dismembrator 250 (Thermo Fisher Scientific, Waltham, MA) under minimal light to prevent photolytic degradation of methylcorrinoids. Unlysed cells were removed by centrifugation at 30,000 × *g* for 1 h at 4 °C. Removal of unlysed cells was confirmed by microscopic examination with a Zeiss Axioscope 2 (Carl Zeiss Microscopy LLC, Thornwood, NY). Total protein concentration of cell lysates was determined by the Bradford method with bovine serum albumin (MilliporeSigma, St. Louis, MO) as standard and the lysates were stored in aliquots at −80 °C in amber glass vials capped with Teflon-lined silicone stoppers and sealed. Cell lysates were used within 2 weeks of preparation.

### Methylation Assays

Methylation assays were performed under strictly anaerobic conditions in the dark at 32 °C inside an anaerobic glove box at a total protein concentration of 1.5 mg/mL for 2 h (standard conditions), unless otherwise specified. To avoid any potential variability that may result from differences in the thiolate concentrations in the cell lysates, the concentration of DTT was kept constant at 2 mM for all experiments. Cell lysates of ND132 WT and *ΔhgcAB* were equilibrated under the desired experimental conditions for 2 minutes before adding a freshly prepared 1.5 μM (300 ppb) HgCl_2_ stock solution to a final concentration of 30 nM (6 ppb), unless otherwise specified. The reactions were stopped at the desired time points by adding 0.5% (v/v) trace-metal grade H_2_SO_4_ (Thermo Fisher Scientific, Waltham, MA), and the samples were moved immediately to a −20 °C freezer and stored until MeHg analysis. All experiments were performed in duplicate. The reaction buffer, PBS, pH 7.0 (without cell lysates) and *ΔhgcAB* cell lysates were used as controls. Samples were processed similarly for total Hg (THg) analysis. Experiment-specific variations used, while keeping all other parameters constant, were as follows. For the time-dependence experiments, aliquots were removed from the reaction mixture before the addition of HgCl_2_ (0 h), and following the addition of HgCl_2_ at 2 min, 30 min, 2, 4, 24, 48, 72 and 92 h. For the pH dependence experiments, the cell lysates were diluted in PBS buffer and solutions were adjusted to pH values of 4.0, 5.0, 6.0, 7.0, 8.0 and 9.0, respectively. For the temperature dependence experiments the samples were preequilibrated before the addition of HgCl_2_ for 5 minutes at the desired temperatures, 4 °C, room temperature (24 °C), 32, 40 and 50 °C, respectively, and following the addition of HgCl_2_ the samples were incubated for 2 h. For the protein concentration dependence experiments, the cell lysates were diluted to final protein concentrations of 0.3, 0.6, 1. 3 and 2.6 mg/mL, respectively. For the substrate concentration dependence experiments, cell lysates were spiked with HgCl_2_ to final concentrations of 0.5, 1, 5, 10, 15, 50 and 60 nM with stock solutions of 50 nM, 0.5 and 1.5 μM, respectively. The effects of pyruvate and CH_3_-H_4_folate were tested by adding a freshly prepared sodium pyruvate stock solution (500 mM) to a final pyruvate concentration of 10 mM and a freshly prepared 10 mg/mL stock solution of CH_3_-H_4_folate (MilliporeSigma, St. Louis, MO) to a final concentration of 16.8 μM. The samples were equilibrated with pyruvate and CH_3_-H_4_folate for 2 minutes before the addition of HgCl_2_. To test the oxygen sensitivity, sets of WT and *ΔhgcAB* cell lysates were incubated at standard conditions inside the glove box (< 0.6 ppm O_2_) and other sets were preequilibrated outside the glovebox (at ambient O_2_) for 2 minutes before the addition of HgCl_2_ and the incubation was continued for 2 h under the same conditions after the addition of HgCl_2_. All other experimental parameters and procedures were followed as described above.

The effect of ATP was tested by adding a freshly prepared stock solution of ATP disodium salt (MilliporeSigma, St. Louis, MO) to a set of vials before the addition of HgCl_2_ (ATP at 0 h) and 1 h after the addition of HgCl_2_ (ATP at 1 h) to achieve final ATP concentrations of 2 mM and 10 mM. The reaction was stopped at 2 h, as described previously. To test the effect of a non-hydrolyzable ATP analog, a freshly prepared stock solution of Adenosine 5’-(β,γ-imido)triphosphate lithium salt hydrate (MilliporeSigma, St. Louis, MO) was added to cell lysates to a final concentration of 5 mM and incubated for 5 minutes before initiating the reaction by addition of HgCl_2_.

### Demethylation assays

Demethylation potentials were measured in the cell lysates of WT and *ΔhgcAB* ND132, the samples were processed under standard conditions described above, except that 5 nM MeHg was used as the substrate instead of Hg(II). A 250 nM MeHg working stock solution was freshly prepared from a 1 ppm methylmercury standard solution (Brooks Rand Instruments, Seattle, WA).

### Total mercury (THg) analysis

All samples were treated with 5% (v/v) BrCl overnight, followed by 5-minute incubation with 30% (w/v) NH_2_OH·HCl before the THg analysis. THg was analyzed by reduction with 0.8% (w/v) stannous chloride (SnCl_2_) on an RA-915+ mercury analyzer (Ohio Lumex, USA). An aliquot of Hg-containing sample was added to an excess of 0.8% (w/v) SnCl_2_ and purged with ultrahigh-purity N_2_. The emerging Hg(0) was quantified by a cold vapor atomic absorption spectroscopy (CV-AAS) Zeeman effect Hg analyzer (Lumex RA-915+, Ohio Lumex Company, Inc. Twinsburg, OH), which was calibrated with a set of Hg standards (Brooks Rand Instruments, Seattle, WA).

### Methylmercury (MeHg) analysis

All samples were processed using a non-aqueous extraction method (45) followed by a modified version of EPA Method 1630 described previously (44). Briefly, Me^200^Hg was added to all the samples as an internal standard, MeHg was then extracted with an acidic KBr solution, followed by extraction of MeHg with CH_2_Cl_2_. After phase separation, CH_2_Cl_2_ was volatilized completely to release MeHg into a pure aqueous phase. MeHg samples were further processed by distillation, ethylation, and trapping on a Tenax column via N_2_-purging. Following thermal desorption and separation by gas chromatography, MeHg was detected by ICP-MS. The recovery of spiked MeHg standards was 100 ± 10%, and the detection limit was about 3·10^-5^ nM MeHg.

### Statistical analysis

GraphPad Prism (GraphPad Software, La Jolla, CA) and the R software environment for statistical computing and graphics (https://www.r-project.org/) were used to analyze and plot the data.

## Acknowledgments

We thank Xiangping Yin and Linduo Zhao at Oak Ridge National Laboratory (ORNL) for technical assistance in mercury and methylmercury analyses. This research was sponsored by the Office of Biological and Environmental Research, Office of Science, U.S. Department of Energy (DOE), as part of the Critical Interfaces Science Focus Area at Oak Ridge National Laboratory, which is managed by UT-Battelle, LLC for the DOE under contract DE-AC05-00OR22725. This manuscript has been authored by UT-Battelle, LLC under Contract No. DE-AC05-00OR22725 with the U.S. Department of Energy.

## Author contributions

S.S.D., J.D.W., S.W.R., J.M.P., and A.J. designed the research. S.S.D. and A.J. performed the experiments and analyses. S.S.D., K.W.R., J.M.P., J.D.W., S.W.R., and A.J. wrote the paper.

