## Supplementary information for "Kinetics of enzymatic mercury methylation at nanomolar concentrations catalyzed by HgcAB"

**This PDF file includes:**

Supplementary experimental procedures
Figs. S1 to S4
Table S1
References for SI reference citations

### Supplementary experimental procedures

#### *Identification of RACo homologs in the genomes of Hg methylators*

The Basic Local Alignment Search Tool (1) was used to search for homologs of the reductive activator of corrinoid protein (RACo) from *Carboxydothermus hydrogenoformans* (locus tag: CHY_1224, UniProt accession number Q3ACS2, annotated as DUF4445 domain protein or [2Fe-2S] ferredoxin) in selected Hg methylators (*Desulfovibrio desulfuricans* ND132, *Methanosphaerula palustris* E1-9c, *Desulfosporosinus youngiae* JW/YJL-B18, *Dethiobacter alkaliphilus* AHT 1, *Desulfotignum phosphitoxidans* FiPS-3 and *Methanoregula boonei* 6A8) from 3 different groups: Proteobacteria, Firmicutes and Archaea (representative of phylogenetic and metabolic diversity of Hg methylators) (E-value ≤ $1\cdot10^{-30}$). The sequence alignment (Fig. S4) was generated using MUSCLE (2).

#### *Free energy of the transition state*

The first-order catalytic rate constant ($k_{\mathrm{cat}}$) describes the limiting rate of an enzyme-catalyzed reaction at saturation. Since the overall rate of the reaction is determined by the intermediate with the highest free energy (the transition state), $k_{\mathrm{cat}}$ is related to the energy of the transition state (3, 4) as follows:

$$k_{cat} = \frac{k_B T}{h} e^{\frac{-\Delta G^{\ddagger}}{RT}}$$

where $k_B$ is the Boltzmann constant, $T$ is the absolute temperature, $h$ is the Planck constant, $R$ is the universal gas constant and $\Delta G^{\ddagger}$ is the relative free energy of the transition state.

Thus, for a given $k_{\mathrm{cat}}$, $\Delta G^{\ddagger}$ can be calculated as follows:

$$\Delta G^{\ddagger} = RT \cdot \ln \frac{k_B T}{k_{cat} h}$$

#### *Estimates for specific activity*

In the absence of studies with purified proteins, we estimate the specific activity of HgcAB based on the following assumptions. The residual amount of MeHg formed in cell lysates exposed to oxygen may be attributable to methylcorrinoids associated with HgcA in the cell lysates prior to oxygen exposure and before the addition of the Hg(II) substrate. Any methylcorrinoids associated with HgcA are not redox-sensitive and should be able to transfer the methyl group to Hg(II), even under aerobic conditions. Once the methyl group is transferred in the presence of oxygen, the corrinoid cannot be reduced back to the Co(I) state. Therefore, the residual amount of MeHg formed after exposure to oxygen and addition of Hg(II) may be the result of a single turnover. We did not observe any significant levels of MeHg in the presence or absence of oxygen in cell lysates of the Δ*hgcAB* mutant, suggesting that Hg methylation by residual methylcorrinoid not associated with HgcA can be ruled out. Assuming that all corrinoid cofactors bound to HgcA are present as methylcorrinoids after exposure to oxygen and prior to the addition of Hg(II), the concentration of MeHg formed in the cell lysates (0.2 nM) should be stoichiometrically equivalent to the concentration of HgcA. Thus, we speculate that the fraction of HgcA is equivalent to approximately 0.0004% (0.2 nM or $7 \cdot 10^{-6}$ mg/mL) of the total protein concentration in cell lysates.

### Supplementary Figures

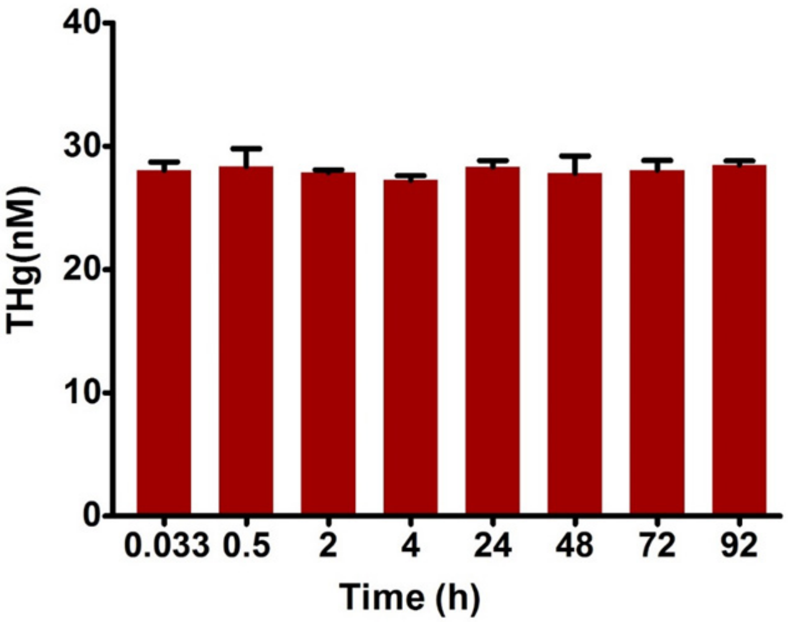


**Fig. S1.** Hg mass balance based on analysis of total Hg (THg) over time. THg concentrations for all samples in the time-dependent Hg methylation studies (**Fig. 2**) were measured as described (N=2).

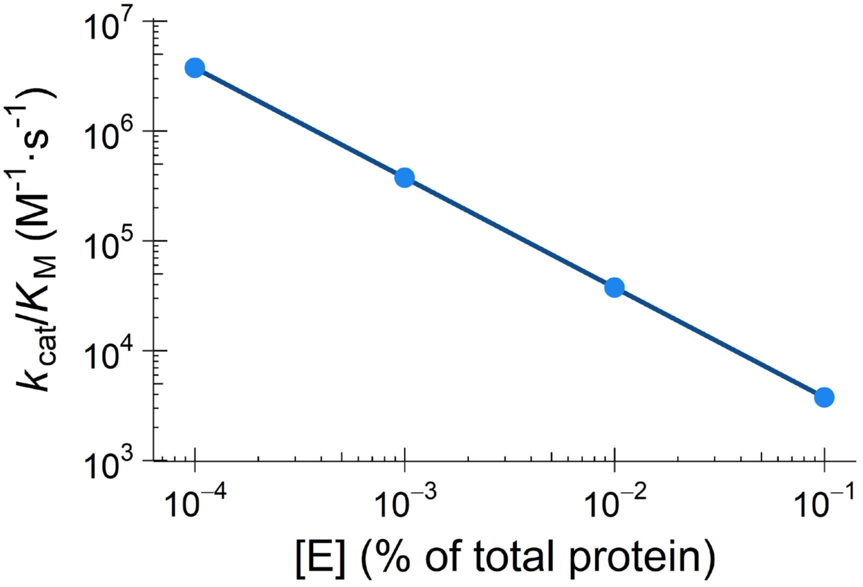


**Fig. S2.** Catalytic efficiency of HgcAB-mediated Hg methylation. Predicted catalytic efficiencies ($k_{cat}/K_M$) of HgcAB-mediated enzymatic Hg methylation for a range of postulated enzyme (HgcA) concentrations expressed as a percentage of total cell protein (**Table 1**).

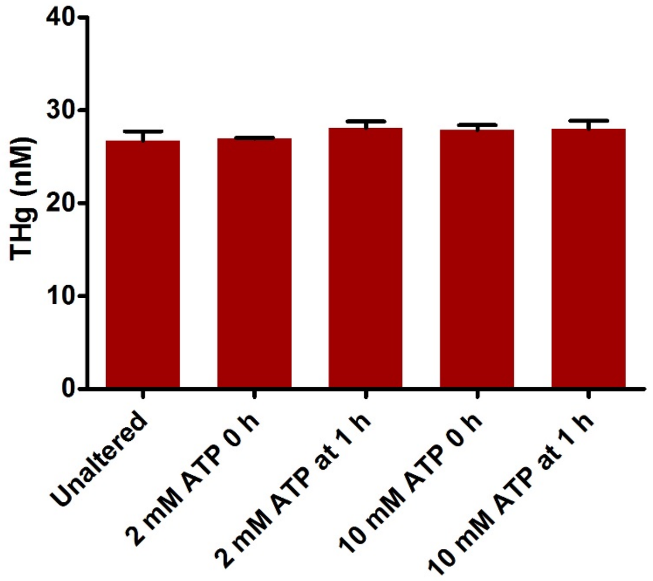


**Fig. S3.** Effect of ATP addition on Hg mass balance. THg concentrations for all samples in the ATP-dependent Hg methylation experiments (Fig. 5C) were determined (N=2).

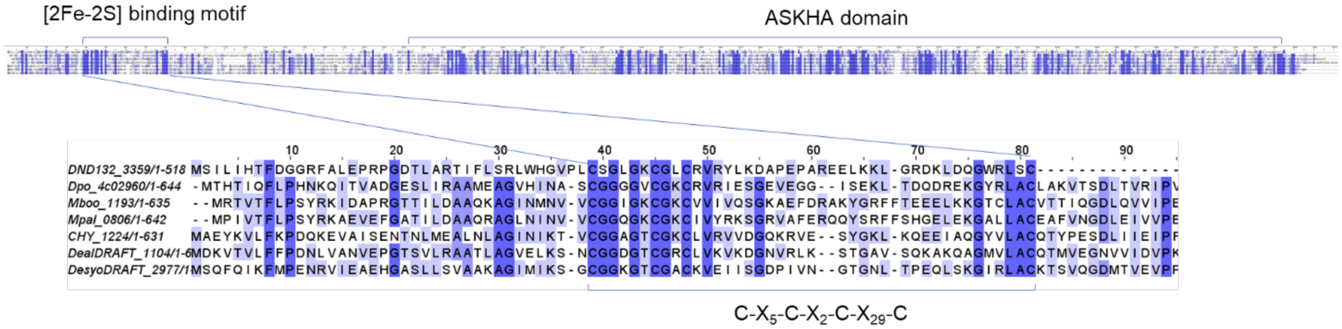


**Fig. S4**. Sequence alignment of genes encoding a reductive activator of corrinoid protein (RACo). MUSCLE-based sequence alignment of RACo from *Carboxydothermus hydrogenoformans* (CHY_1224) and selected homologs from Hg methylators (**Table S1**). Conserved residues are indicated in blue, with the shade of blue corresponding to the level of conservation (darker = higher). RACo harbors one [2Fe-2S] binding site and an acetate and sugar kinase/heat shock cognate/actin (ASKHA) domain.

**Supplementary Tables**

**Table S1:** Homologs of RACo in the genomes of selected Hg methylators.

| **Hg Methylator** | **Group** | **Locus tag (HgcA, HgcB)** | **RACo locus tag** | **% Identity to RACo (CHY_1224)** |
|---|---|---|---|---|
| *Desulfovibrio desulfuricans* ND132 | Proteobacteria | DND132_1056, DND132_1057 | DND132_3359 | 23.88 |
| *Desulfotignum phosphitoxidans* FiPS-3 | Proteobacteria | Dpo_8c00130, Dpo_8c00140 | Dpo_4c02960 | 40.64 |
| *Desulfosporosinus youngiae* DSM 17734 | Firmicutes | DesyoDRAFT_4238, DesyoDRAFT_4237 | DesyoDRAFT_4022 | 48.68 |
| *Dethiobacter alkaliphilus* AHT 1 | Firmicutes | DealDRAFT_3158, DealDRAFT_3157 | DealDRAFT_1104 | 54.99 |
| *Methanoregula boonei 6A8* | Archaea | Mboo_0422, Mboo_0421 | Mboo_1193 | 39.62 |
| *Methanosphaerula palustris* E1-9c | Archaea | Mpal_1034, Mpal_1035 | Mpal_0806 | 39.09 |
